# MicroRNA-dependent inhibition of PFN2 orchestrates ERK activation and pluripotent state transitions by regulating endocytosis

**DOI:** 10.1101/2020.02.06.936070

**Authors:** Carolyn Sangokoya, Robert Blelloch

## Abstract

Profilin2 (PFN2) is a target of the embryonic stem cell (ESC) enriched miR-290 family of miRNAs and an actin/dynamin binding protein implicated in endocytosis. Here, we show that the miR-290-PFN2 pathway regulates many aspects of ESC biology. In the absence of microRNAs, PFN2 is upregulated in ESCs, with a resulting decrease in endocytosis. Reintroduction of miR-290, knockout of PFN2, or disruption of the PFN2 dynamin interacting domain in miRNA deficient cells reverses the endocytosis defect. The loss of miRNA suppression of PFN2 and associated reduction in endocytosis impairs ERK signaling, which in turn inhibits ESC cell cycle progression and differentiation from a naïve to formative state. Mutagenesis of the single canonical conserved 3’UTR miR-290 binding site of PFN2 in otherwise wild-type cells recapitulates these phenotypes. Together, these findings define an axis of post-transcriptional control, endocytosis, and signal transduction that is essential for ESC self-renewal and differentiation.

---

MicroRNAs (miRNAs) arising from the miR-290 family are highly enriched in pluripotent stem cells^1–4^ and are active over much of early mammalian embryonic development^5,6^. The miR-290 miRNA family also enhances the promotion of somatic cell reprogramming to induced pluripotency^7–9^. A number of confirmed miR-290 targets are preferentially suppressed in pluripotency, and upon knockdown also enhance somatic cell reprogramming to induced pluripotency^7,8^. One of these targets, Profilin2 (PFN2), is an evolutionarily highly conserved regulator of actin cytoskeletal dynamics^10^ and has been shown to be a mediator of endocytosis^11–13^. How PFN2 regulates pluripotency is unknown.

To confirm miRNA-mediated repression of PFN2 in pluripotent stem cells we first examined endogenous PFN2 in wild-type versus miRNA-deficient *Dgcr8*-KO mouse embryonic stem cells (ESCs) and found significantly increased PFN2 in *Dgcr8*-KO ESCs at the protein and mRNA levels (Fig. 1a-b). Addback of the miR-290 family member miR-294 into *Dgcr8*-KO ESCs by mimic transfection returned PFN2 expression to wild-type levels, while addback of miR-294 mimic with a mutant seed sequence did not (Fig. 1b).

**Figure 1.**
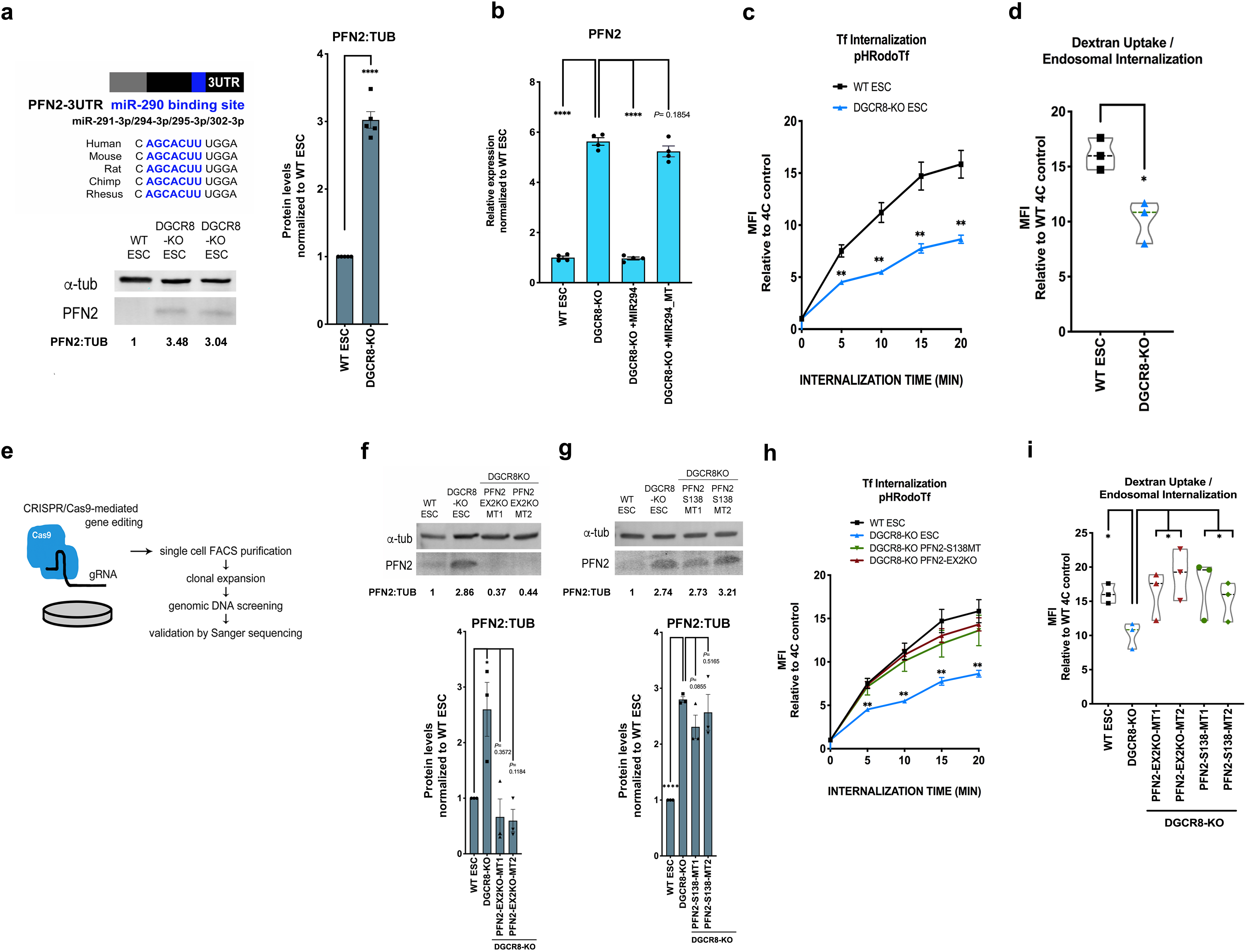
microRNAs regulate endocytosis through their inhibition of PFN2 in pluripotent stem cells. (a) Upper left panel, schematic depicting the PFN2 3’UTR with conserved miR-290 binding site highlighted in blue. Lower left panel, western blot analysis of endogenous PFN2 in v6.5 ESC (WT) and *Dgcr8*-KO ESC. Representative image is shown. Right panel, summary quantification of five independent westerns. Error bars, SEM. \*\*\*\**P* < 0.0001 (unpaired two-tailed t-test). UTR, untranslated region; ESC, embryonic stem cells. (b) qRT-PCR analysis of PFN2 expression in wild-type (WT), *Dgcr8*-KO, *Dgcr8*-KO with addback of miR-294 mimic (+MIR294) or miR-294 mimic with mutant seed (MIR294_MT). n=4 independent experiments. Error bars represent s.e.m. \*\*\*\**P* < 0.0001 (unpaired two-tailed t-test). (c) pH-labeled transferrin endosomal uptake assay in wild-type (WT) and *Dgcr8*-KO ESCs over indicated times, measured by change in mean fluorescence intensity (MFI) relative to control. *n* = 3 independent experiments. Error bars represent s.e.m. \*\**P* < 0.01 (unpaired two-tailed t-test). (d) pH-labeled dextran endosomal uptake assay in wild-type (WT) and *Dgcr8*-KO ESCs, measured by change in mean fluorescence intensity (MFI) relative to control. *n* = 3 independent experiments. Data shown as median ± quartiles. \**P* < 0.05 (unpaired two-tailed t-test). (e) Schematic representation of CRISPR/Cas9-mediated gene editing strategy to generate PFN2 mutant ESC lines. (f) Top panel, western blot analysis of PFN2 expression in wild type (WT) ESC, *Dgcr8*-KO, and two clones (MT1 and MT2) of PFN2-EX2KO mutant ESCs. Representative image is shown. Bottom panel, Summary quantification of western blot data. n= at least 3 independent experiments. (g) Top panel, western blot analysis of PFN2 expression in wild type (WT) ESC, *Dgcr8*-KO, and two clones (MT1 and MT2) of PFN2-S138 mutant ESCs. Representative image is shown. Bottom panel, Summary quantification of western blot data. n= at least 3 independent experiments. (h) pH-labeled transferrin endosomal uptake assay in wild-type (WT), *Dgcr8*-KO, PFN2-EX2KO, and PFN2-S138 ESCs over indicated times, measured by change in mean fluorescence intensity (MFI) relative to control. *n* = 3 independent experiments. Error bars represent s.e.m. \*\**P* < 0.01 (unpaired two-tailed t-test). (i) pH-labeled dextran endosomal uptake assay in wild-type (WT), *Dgcr8*-KO, PFN2-EX2KO, and PFN2-S138 ESCs measured by change in mean fluorescence intensity (MFI) relative to control. *n* = 3 independent experiments. data shown as median ± quartiles. \**P* < 0.05 (unpaired two-tailed t-test). Scanned images of unprocessed blots are shown in Supplementary Fig. 5.

Given PFN2’s reported role in endocytosis, we next asked whether miRNAs are required for normal endocytosis in ESCs. To test this, a canonical assay for receptor-mediated endocytosis, uptake of a pH-sensitive labeled transferrin, was performed in wild-type versus *Dgcr8*-KO ESCs (Fig. 1c). *Dgcr8*-KO ESCs demonstrated significantly decreased transferrin uptake. Addback of miR-294 mimic increased transferrin uptake to wild-type levels (Supplementary Fig. 1a). To study non-receptor-mediated dynamin-dependent endocytosis, endosomal uptake of pH-sensitive labeled nonspecific dextran was also performed and *Dgcr8*-KO ESCs demonstrated significantly decreased dextran uptake (Fig. 1d). These data show that a miRNA-dependent mechanism regulates endocytosis in pluripotent stem cells.

PFN2 has been shown to interact directly with the proline-rich domain of Dynamin^11^ and inhibit endocytosis in neuronal and non-neuronal contexts^11–12,14^. Previous studies show that the Dynamin proline-rich domain is important for co-localization with clathrin, as deletion of the entire proline-rich region abolishes co-localization of dynamin with clathrin^15^. Importantly, disruption of PFN2 at S138 abrogates the PFN2:Dynamin interaction and abrogates inhibition of endocytosis^11^. To study the potential role for PFN2 upregulation in the endocytosis defect seen with miRNA loss, we used CRISPR-genomic editing to make both a total PFN2 knockout and a specific mutation to the S138 region in the *Dgcr8*-null background. To produce the *Pfn2* knockout, the second exon was disrupted (PFN2ΔEX2, Fig. 1e, Supplementary Fig. 1b) resulting in a dramatic reduction in mRNA and loss of detectable protein by western (Fig. 1f). In contrast, specific mutagenesis of the PFN2 S138 region (Supplementary Fig. 1c) resulted in wild-type mRNA and protein levels (Fig. 1g, Supplementary Fig. 1d). These *Dgcr8*; *Pfn2* double mutants were then evaluated for changes in endocytosis. Both receptor and non-receptor mediated endocytosis were significantly enhanced in the double mutants relative to *Dgcr8*-KO alone, reaching levels similar to wild-type cells (Fig. 1h-i). These data suggest that upregulation of PFN2 alone could account for the much of the endocytosis defect seen in the miRNA deficient cells. Furthermore, the interaction of PFN2 with dynamin is critical for its ability to suppress endocytosis in ESCs.

Next we sought to determine the significance of PFN2-regulated endocytosis in pluripotent stem cell biology. Since the miR-290 family miRNAs play a role in the proliferation^16–18^, unique cell-cycle structure^19^, and differentiation of ESCs ^16,18^, we looked at the potential contribution of PFN2 levels in each of these essential miRNA-mediated processes. As previously described, the loss of Dgcr8 leads to a decrease in the growth rate of ESCs (Fig. 2a). The loss of PFN2 protein or the PFN2:Dynamin interaction domain in the *Dgcr8* KO background largely rescued this defect (Fig. 2a). The addition of a miR-294 mimic showed a further, albeit small, increase in the rate suggesting additional targets for the microRNA in growth control (Fig 2b). Also, as previously described, miRNA-deficient *Dgcr8*-KO ESCs showed an accumulation of cells in G1 relative to that seen in wild-type cells^19,20^ (Fig. 2c-d). Loss of PFN2 protein or the PFN2:Dynamin interaction in the miRNA-deficient *Dgcr8*-KO ESCs shifted the cell cycle structure to that much more similar to wild-type ESCs (Fig. 2e-g). Addback of miR-294 in the *Dgcr8*; *Pfn2* double knockouts showed a further, albeit slight, correction in the cell cycle structure of the cells (Fig. 2h-l). These data show that the loss of PFN2 function can in large part reverse the cell growth and cell cycle defects seen with the loss of miRNAs.

**Figure 2.**
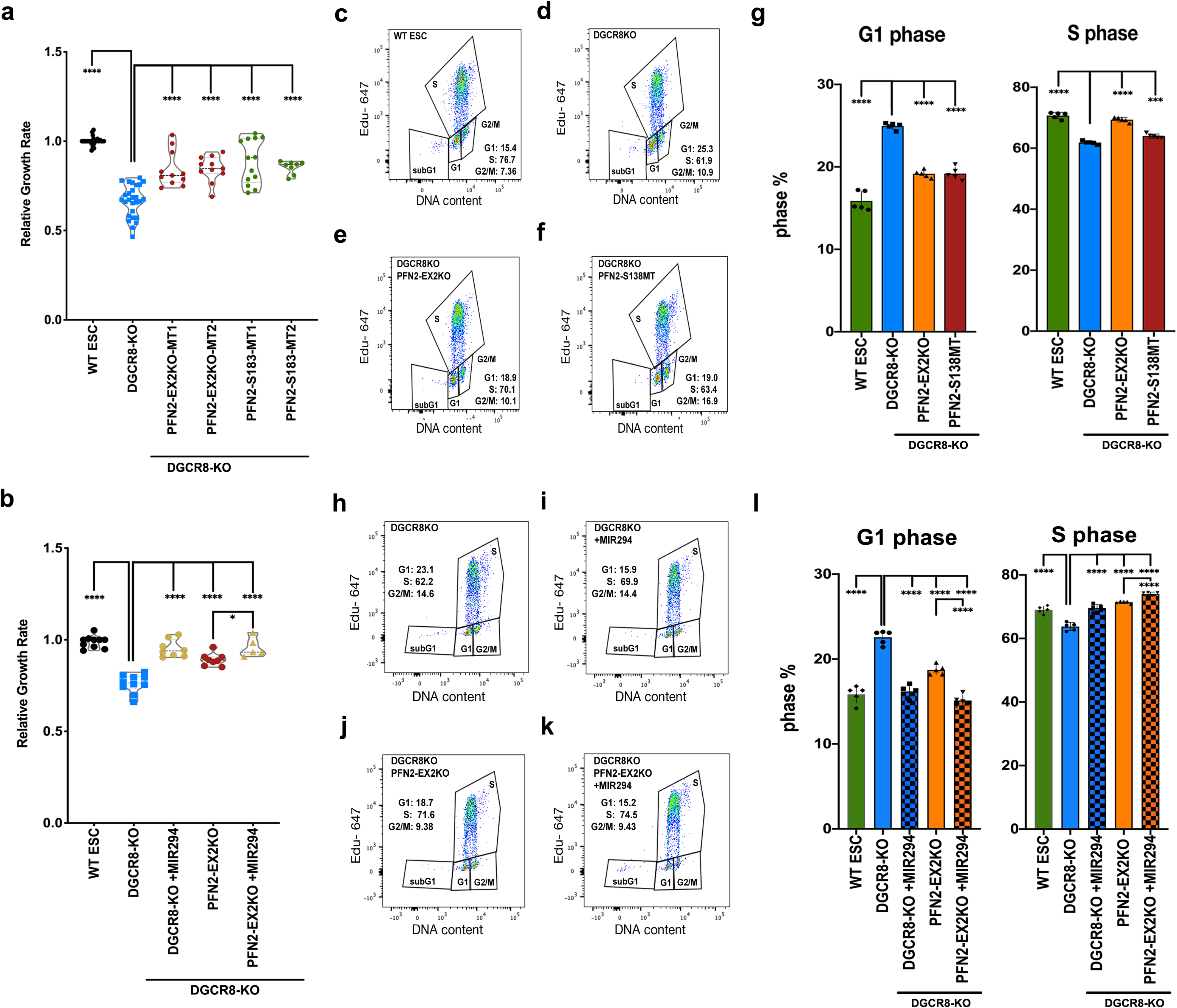
Elevated PFN2 in miRNA deficient cells suppresses growth rate and inhibits G1/S transition in ESCs. (a) Growth rates of indicated ESCs relative to wild-type (WT) ESCs. *n* = at least 8 independent experiments. data shown as median ± quartiles. \*\*\*\**P* < 0.0001 (unpaired two-tailed t-test). (b) Growth rate of indicated ESCs relative to wild-type (WT) ESCs. *n* = at least 5 independent experiments. data shown as median ± quartiles. \*\*\*\**P* < 0.0001 (unpaired two-tailed t-test). Error bars represent s.e.m. \*\*\*\**P* < 0.0001, \*\*\**P* < 0.001 (unpaired two-tailed t-test). (c-f) Representative dot plot depicting dual Click-IT-based EdU and FxCycle DNA content analysis of indicated ESC sample from n=5 independent experiments (g) Summary statistics of G1 phase distribution (left panel) and S-phase distribution (right panel) from n=5 independent experiments. (h-k) Representative dot plot depicting dual Click-IT-based EdU and FxCycle DNA content analysis of indicated ESC sample from n=5 independent experiments (l) Summary statistics of G1 phase distribution (left panel) and S-phase distribution (right panel) from n=5 independent experiments. Error bars represent s.e.m. \*\*\*\**P* < 0.0001 (unpaired two-tailed t-test).

In pluripotent stem cells, the MAPK/ERK signaling pathway is important in regulation of stem cell growth and cell cycle progression in serum+LIF^21–22^. Therefore, we asked whether a miRNA-PFN2 axis regulates ERK signaling in ESCs. ERK is activated by phosphorylation^23^. Basal phosphorylated ERK (pERK) levels were decreased in *Dgcr8*-KO ESCs compared to wild-type (Fig. 3a). This decrease was largely reversed with the simultaneous loss of PFN2 and partially reversed with deletion of the PFN2-Dynamin interaction domain (Fig. 3b-c). Next, we asked whether this role for the miRNA-PFN2 axis in ERK signaling could explain the proliferation defect associated with *Dgcr8* loss. The MEK inhibitor PD0325901 suppressed growth in all the genetic backgrounds: *Dgcr8*-KO, and PFN2-mutant *Dgcr8*-KO ESCs cells (Fig. 3d). Importantly, however, the positive impact of the knockout of PFN2 or the mutagenesis of PFN2’s interaction domain with Dynamin on the growth of *Dgcr8* null cells was lost in the presence of the MEK inhibitor. To capture the range of in vitro pluripotency conditions we additionally cultured the pluripotent stem cells with GSK3 inhibition (CHIR99021) with and without added MEK inhibition and performed growth rate studies. Again, inhibition of ERK activation extinguished the increased growth rate seen in PFN2-mutant *Dgcr8*-KO ESCs (Supplementary Fig. 2a). These results indicate that ERK-dependent signaling acts downstream of PFN2-regulated endocytosis in the rapid growth rate of ESCs.

**Figure 3.**
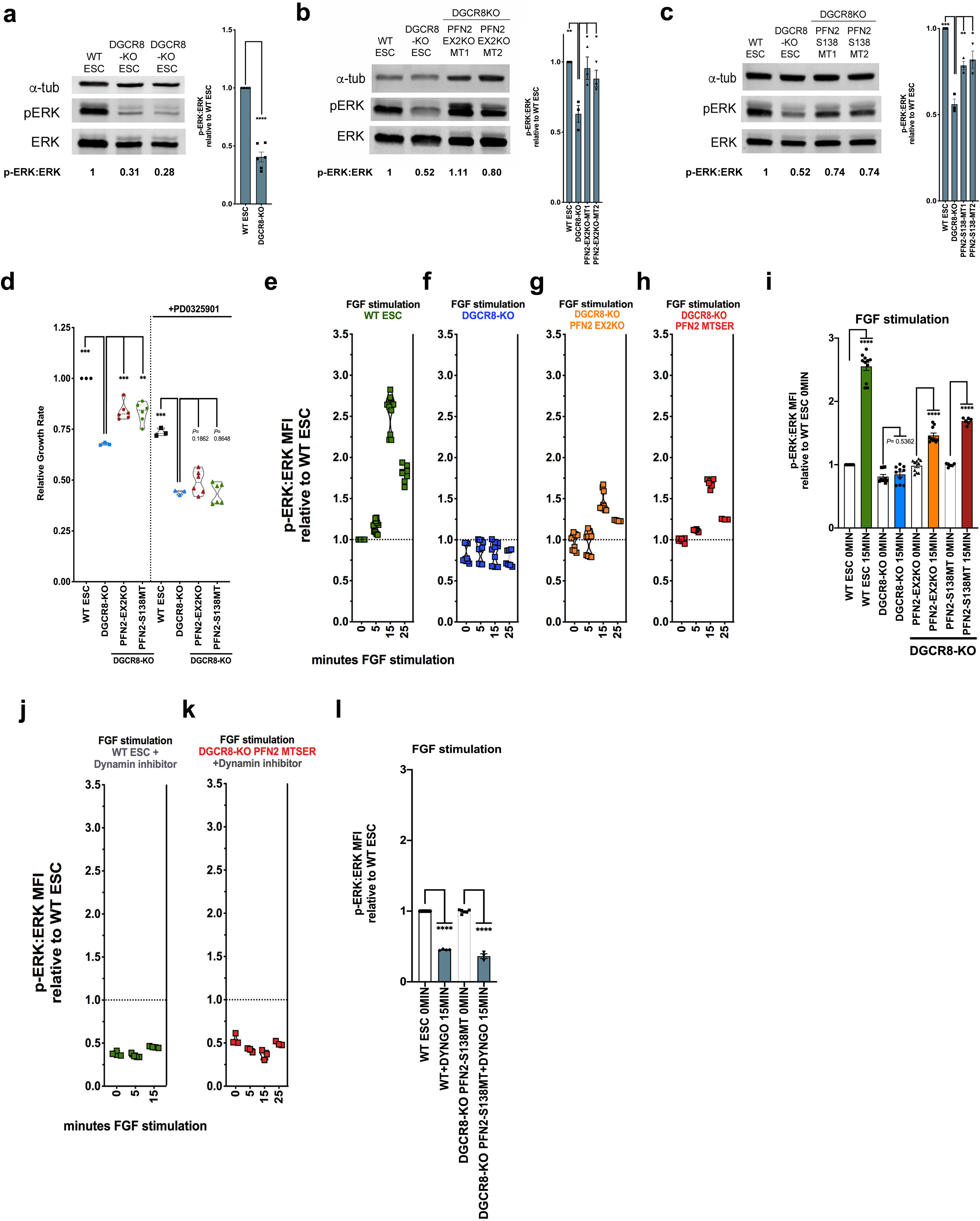
Elevated PFN2 in miRNA deficient ESCs regulates ERK activity in dynamin-dependent manner. (a) Left panel, western blot analysis of ERK and phospho-ERK expression in wild type (WT) ESC and *Dgcr8*-KO ESCs. Representative image is shown. Right panel, Summary quantification of western blot data of at least 3 independent experiments. (b) Left panel, western blot analysis of ERK and phospho-ERK expression in wild type (WT) ESC, *Dgcr8*-KO, and two clones (MT1 and MT2) of PFN2-EX2; Dgcr8 DKO mutant ESCs. Representative image is shown. Right panel, Summary quantification of western blot data of at least 3 independent experiments. (c) Left panel, western blot analysis of ERK and phospho-ERK expression in wild type (WT) ESC, *Dgcr8*-KO, and two clones (MT1 and MT2) of PFN2-S138; Dgcr8 KO mutant ESCs. Representative image is shown. Right panel, Summary quantification of western blot data of at least 3 independent experiments. (d) Growth rates of indicated ESCs relative to wild-type (WT) ESCs in presence or absence of MEK inhibitor, PD0325901. *n* = 3 independent experiments. Data shown as median ± quartiles. \*\*\**P* < 0.001, \*\**P* < 0.01 (unpaired two-tailed t-test). (e-h) Ratio of intracellular phospho-ERK to ERK protein levels in ESCs of indicated genetic background and at indicated time following FGF4 stimulation relative to wild-type (WT) at time 0. *n* = 3 independent experiments. Data shown as median ± quartiles. (i) Summary statistics from FGF stimulation assay at min (0 min) and max (15 min) timepoints shown in (e-h) \*\*\*\**P* < 0.0001 (unpaired two-tailed t-test). (j-k) FGF stimulation assay ratio of intracellular phospho-ERK to ERK protein levels from indicated ESC sample with addition of the dynamin inhibitor Dyngo4a, relative to wild-type (WT) ESC basal levels over indicated time course. *n* = 3 independent experiments. Data shown as median ± quartiles. (l) Summary statistics from FGF stimulation assay at 0 and 15 minute timepoints shown in (j-k) \*\*\*\**P* < 0.0001 (unpaired two-tailed t-test). Scanned images of unprocessed blots are shown in Supplementary Fig. 5.

The MAPK/ERK signaling pathway is an important regulator in the transition from pluripotency to early lineage commitment^24^. In pluripotency, LIF overrides the autoinductive capacity of fibroblast growth factor (FGF) –induced MAPK/ERK pathway activation necessary for exiting pluripotency^24^. To directly interrogate the effect of PFN2 on ERK activation under serum+LIF conditions, we performed FGF stimulation assays in wild-type, *Dgcr8*-KO, and PFN2-mutant *Dgcr8*-KO ESCs at single cell resolution. To achieve this, the cells were subjected to short-term FGF4-stimulation, a physiologically relevant cue that induces MAPK/ERK signaling and ERK activation in PSCs^24–25^, and ERK activation was measured by intracellular staining and flow cytometry. As expected, ERK activation was induced in wild-type ESCs (Fig. 3e). However, this induction was absent in *Dgcr8*-KO ESCs (Fig. 3f and Supplementary Fig. 2b-c). Knockout for PFN2 or mutation of its dynamin interaction domain in *Dgcr8*-KO ESCs partially rescued ERK activation (Fig. 3g-i and Supplementary Fig. 3b-d). To further evaluate the role of dynamin-dependent endocytosis in FGF signaling, we treated cells with the dynamin inhibitor Dyngo-4a. As expected, the inhibitor blocked receptor-mediated endocytosis (Supplementary Fig. 2e). Furthermore, the inhibitor reduced basal ERK activity by half and blocked FGF stimulation of phospho-ERK levels in wild-type cells (Fig. 3j). Mutation of the PFN2 dynamin interaction domain did not rescue the effect of this inhibitor (Fig. 3k-l). These data place dynamin-mediated endocytosis downstream of PFN2 and upstream of FGF-mediated ERK activation.

In addition to regulating ESC proliferation, the miR-290 microRNA family is required for the transition of naïve ESCs to post-implantation epiblast-like cells (EpiLC)^26^. This transition also requires a pulse in ERK activation^24–25^. Therefore, we wondered whether the upregulation of PFN2 in *Dgcr8* knockout cells could in part explain their differentiation defect. ESCs were grown in serum+LIF/2i to mimic the naïve state followed by LIF/2i removal, allowing for induction of ERK activation in the absence of LIF. These conditions induce a transition to EpiLCs in 48-72 hours^27–28^. Indeed, wild-type ESCs downregulated naïve markers (*Esrrb*, *Prdm14*, *Rex1*) and upregulated EpiLC markers (*Otx2*, *Dnmt3b*) over a 48 hour time course (Figure 4a,b, Supplementary Fig. 3a). In contrast, however, *Dgcr8*-KO cells showed a dramatic reduction in both the downregulation of the naïve markers and upregulation of EpiLC markers. The introduction of miR-294 mimic into the *Dgcr8*-KO cells largely rescued the defect (Supplementary Fig. 3a-b). Loss of PFN2 also rescued the defect, although to a slightly smaller degree (Figure 4a,b, Supplementary Fig. 3a). These data suggest that the downregulation of PFN2 by miR-290 family of miRNAs is critical for the differentiation of naïve ESCs to EpiLCs, presumably by a enabling a pulse in ERK activation which in turn is regulated by receptor-mediated endocytosis.

**Figure 4.**
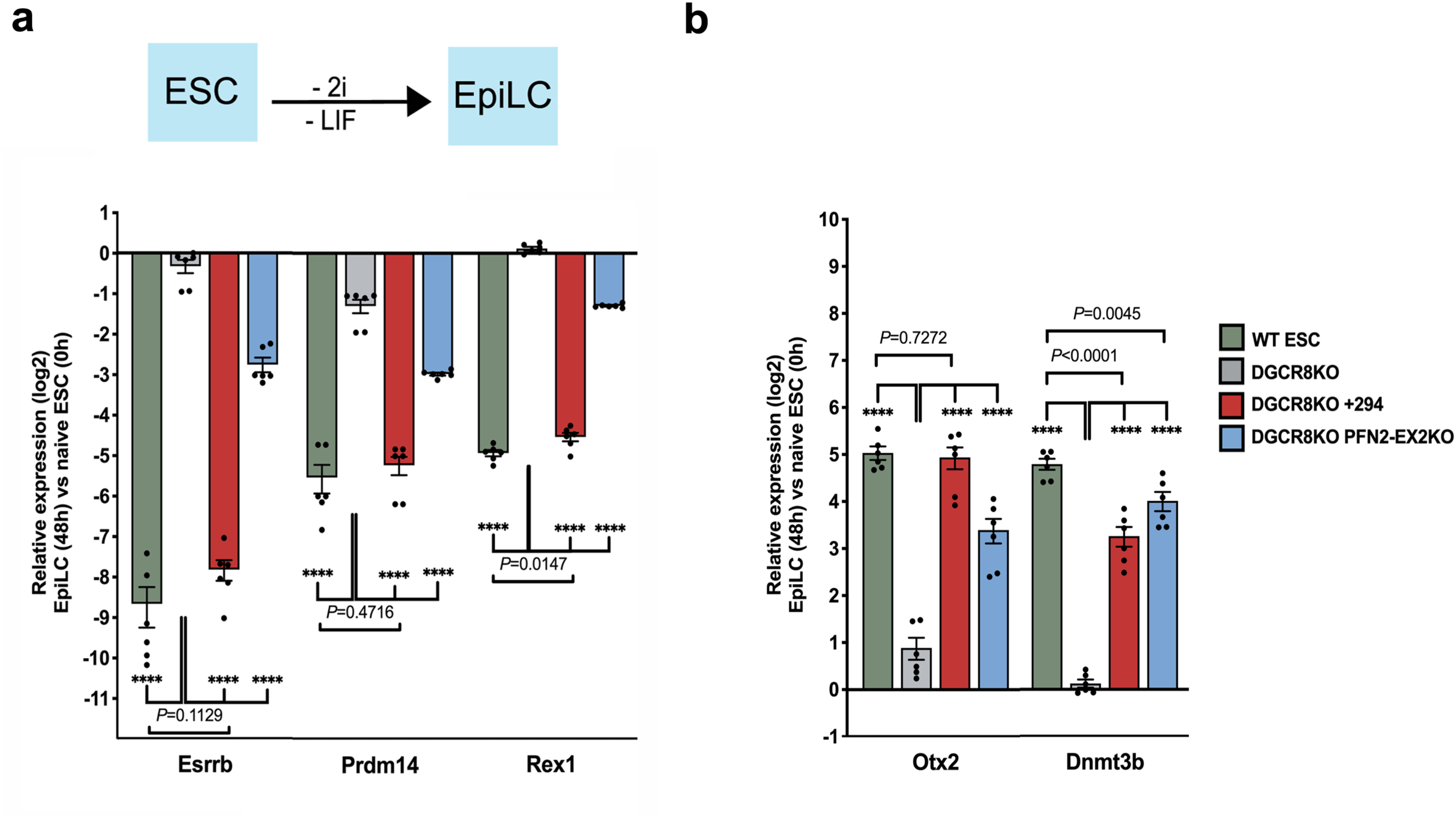
miR-294 addback and PFN2 depletion rescues pluripotency transition defects in *Dgcr8*-KO ESCs. (a) Top panel, schematic ESC to EpiLC transition strategy. Bottom panel, qRT-PCR analysis of naive factors *Esrrb*, *Prdm14*, and *Rex1* expression after transition to EpiLC (48h) relative to naïve state (0h) in wild-type (WT), *Dgcr8*-KO, *Dgcr8*-KO with addback of miR-294 mimic (+294) or *Dgcr8*; *PFN2*-DKO (PFN2-EX2KO) ESCs. *n* = 3 independent experiments. Error bars represent s.e.m. \*\*\*\**P* < 0.0001 (unpaired two-tailed t-test). (b) qRT-PCR analysis of EpiLC markers *Otx2* and *Dnmt3b* expression after transition to EpiLC (48h) relative to naïve state (0h) in wild-type (WT), *Dgcr8*-KO, *Dgcr8*-KO with addback of miR-294 mimic (+294) or *Dgcr8*; *PFN2*-DKO (PFN2-EX2KO) ESCs *n* = 3 independent experiments. Error bars represent s.e.m. \*\*\*\**P* < 0.0001 (unpaired two-tailed t-test). Scanned images of unprocessed blots are shown in Supplementary Fig. 5.

As PFN2 has one canonical miR-290 target site in its 3’UTR^29^, we sought to characterize the singular effect of this site on the phenotypes described above. Using CRISPR-mediated genomic editing, we mutated the region of the PFN2 3’UTR that complements the seed sequence for the miR-290 family (Fig. 5a). These WT-PFN2-3UTRΔ290 mutant ES cells showed elevated PFN2 RNA levels relative to wild-type ES cells, although not to the same degree as the loss of *Dgcr8* (compare Fig. 5a to Fig. 1b). Still, the mutation of this target site led to decreased receptor and non-receptor mediated endocytosis (Fig. 5b,c), decreased growth rate, increased fraction of cells in G1 phase in serum+LIF conditions (Fig. 5d-g), and decreased basal phosphorylated ERK (pERK) (Fig. 5h). To evaluate the impact of the mutant PFN2Δ290 3’UTR on activation of FGF/ERK signaling with ESC differentiation, we measured ERK activation for the first three hours following removal of LIF and 2i from cells grown in naïve culture conditions, matching the time window of the reported pulse of ERK activation during naïve to formative transition^25^. Wild-type cells showed a peak in activation between 30-60 minutes (Fig. 5i, Supplementary Fig. 4). PFN2-3UTRΔ290 mutant cells showed a similar pulse in activation at 30 minutes, but were unable to sustain activation at 60 minutes. This defect was associated with a decrease in activation of differentiation markers at 48 hours post-differentiation (Fig. 5j), while the ability to downregulate naïve factors was not significantly changed (Fig. 5k). Together, these data uncover an important role for the canonical miR-290 target site in the regulation of PFN2 levels and associated downstream phenotypes.

**Figure 5.**
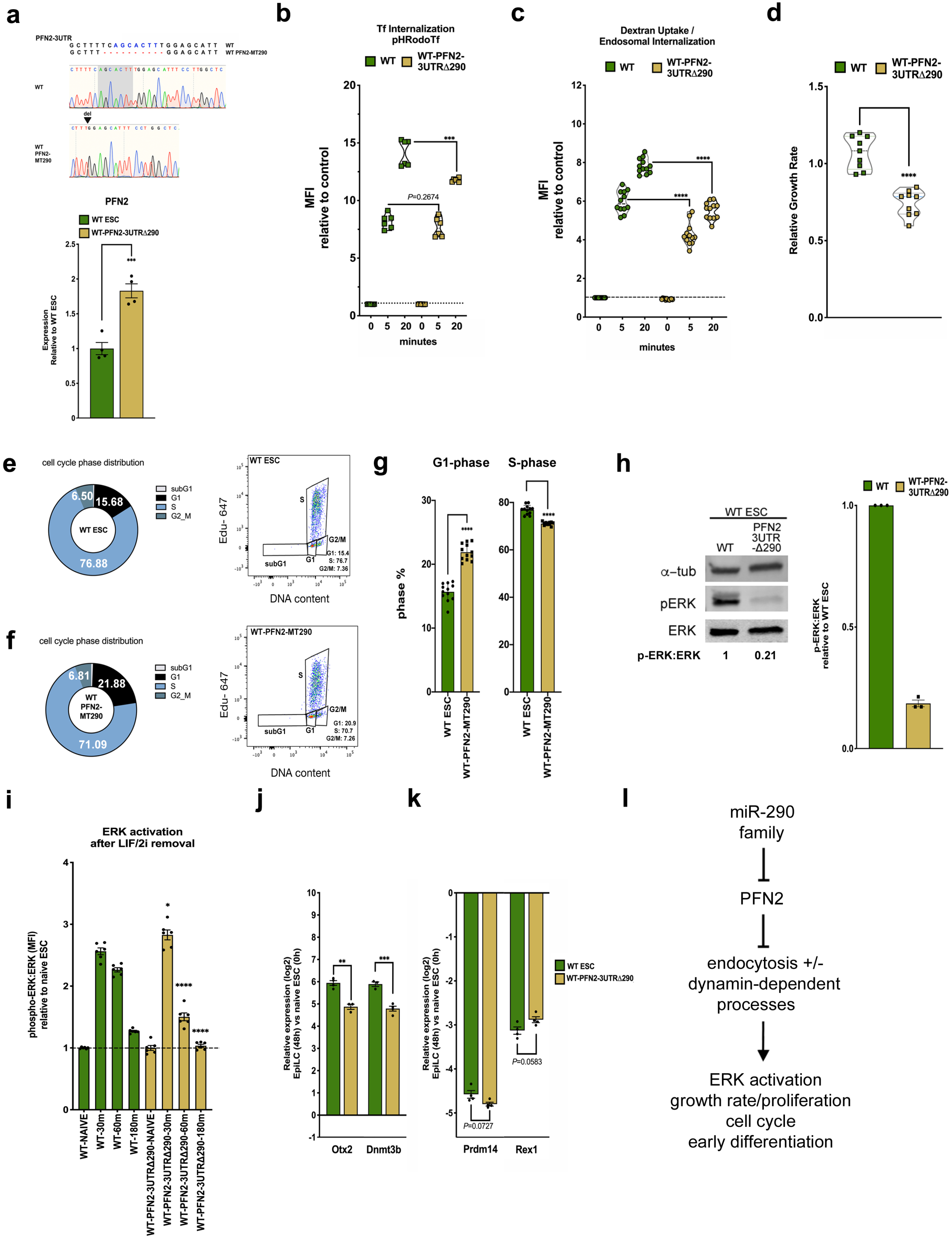
Deletion of single miR-290 seed site in ESCs recapitulates defects seen in *Dgcr8*-KO ESCs. (a) Top panel, Sanger sequencing results showing CRISPR induced mutations to PFN2-3UTR miR-290 seed site in wild-type ESCs, Bottom panel, qRT-PCR analysis of PFN2 expression in wild-type (WT) and WT-PFN2-3UTRΔ290 mutant ESCs. *n* = 4 independent experiments. Error bars represent s.e.m. *** *P* < 0.001 (unpaired two-tailed t-test). (b) pH-labeled transferrin endosomal uptake assay in wild-type (WT) and WT-PFN2-3UTRΔ290 mutant ESCs over indicated times, measured by change in mean fluorescence intensity (MFI) relative to control. *n* = 3 independent experiments. Error bars represent s.e.m. *** *P* < 0.001 (unpaired two-tailed t-test). (c) pH-labeled dextran endosomal uptake assay in wild-type (WT) and WT-PFN2-3UTRΔ290 mutant ESCs, measured by change in mean fluorescence intensity (MFI) relative to control. *n* = 3 independent experiments. Data shown as median ± quartiles. \*\*\*\**P* < 0.0001 (unpaired two-tailed t-test). (d) Growth rates of indicated WT-PFN2-3UTRΔ290 mutant ESCs relative to wild-type (WT) ESCs under indicated conditions. *n* = 3 independent experiments. Data shown as median ± quartiles. \*\*\*\**P* < 0.0001 (unpaired two-tailed t-test). (e-f) Left panel, mean cell cycle phase distribution of indicated ESC sample. Right panel, Representative dot plot depicting dual Click-IT-based EdU and FxCycle DNA content analysis of indicated ESC sample from *n* = 3 independent experiments. (g) Summary statistics of G1 phase distribution (left panel) and S-phase distribution (right panel) from *n* = 3 independent experiments. Error bars represent s.e.m. \*\*\*\**P* < 0.0001 (unpaired two-tailed t-test). (h) Intracellular phospho-ERK protein levels relative ERK protein levels in wild-type (WT) ESC and WT-PFN2-3UTRΔ290 mutant ESC samples relative to naïve conditions over indicated time course. *n* = 3 independent experiments. Data shown as median ± quartiles. Error bars represent s.e.m. \**P* < 0.05, \*\*\*\**P* < 0.0001 (unpaired two-tailed t-test). (i) qRT-PCR analysis of EpiLC markers *Otx2* and *Dnmt3b* after transition to EpiLC (48h) relative to naïve state (0h) in wild-type (WT) and WT-PFN2-3UTRΔ290 mutant ESCs *n* = 3 independent experiments. Error bars represent s.e.m. \*\**P* < 0.01, *** *P* < 0.001 (unpaired two-tailed t-test). (j) qRT-PCR analysis of naive factors *Prdm14*, and *Rex1* expression after transition to EpiLC (48h) relative to naïve state (0h) in wild-type (WT) and WT-PFN2-3UTRΔ290 mutant ESCs *n* = 3 independent experiments. Error bars represent s.e.m. unpaired two-tailed t-test. (k) Summary model for effect of miR-294-dependent target PFN2 effect in pluripotent stem cell biology. Scanned images of unprocessed blots are shown in Supplementary Fig. 5.

Our findings uncover a crucial role for miRNA repression of PFN2 in fine tuning the levels of endocytosis in ESCs which in turn is essential for normal FGF/ERK signaling, ESC self-renewal, and differentiation (Fig. 5l). We show that both basal and FGF-induced ERK activation in ESCs are dependent on dynamin-regulated endocytosis. It has long been proposed that endosomal membranes may serve as platforms for the formation of functional MAPK/ERK pathway signaling complexes^30–33^. While the most comprehensive studies have centered almost exclusively on the epidermal growth factor (EGF)/EGF receptor-stimulated MAPK/ERK pathway^34–35^, the ERK pathway in undifferentiated ES cells is essential and episodically activated predominantly by autocrine signaling by FGF4 through FGF receptors^24^, thus offering an ideal model to re-examine this hypothesis. Intriguingly, clathrin-mediated dynamin-dependent endocytosis has been reported to play a crucial role in maintaining a fine balance between antagonistic signaling pathways in ESC pluripotency, suggesting an additional layer of regulation for the pluripotent state^36^. Future studies examining the spatiotemporal localization of the PFN2:Dynamin:FGF receptor relationship in relation to ERK activation over the early phases of ESC to EpiLC transition could help to further clarify the possibility of a contributory endosomal signaling cascade and other dynamin-dependent processes in pluripotent state transitions.

The results of this study demonstrate clear biologic effects of the post-transcriptional regulation of PFN2 (reduced growth rate, reduced endocytosis, reduced ERK activation, altered cell-cycle structure) following removal of its single canonical miR-290 site. Yet data from microRNA-deficient ESCs (*Dgcr8* loss) with add-back of miR-294 demonstrates a more robust effect on PFN2 expression and associated downstream phenotypes, suggesting that 1) other microRNA-mediated sites of regulation in the PFN2 3UTR, including possible non-canonical miR-290 sites, likely coordinately control PFN2 mRNA stability and that 2) other microRNA-mediated targets may secondarily affect PFN2 levels. Further determination of the precise mechanisms by which other miR-290-regulated targets contribute to the overall effect of endocytic mechanisms, cell-cycle structure, cell growth rates, and ERK activation, as well as PFN2 levels, will continue to be an important goal in dissection of the molecular and signaling dynamics underlying pluripotency and pluripotent state transitions.

The fine tuning of endocytosis and the resulting impact on cell fate decisions is a relatively unexplored area. Here we show that miR-290’s fine tuning of endocytosis is crucial for enabling normal levels ERK signaling during differentiation of embryonic stem cells toward the formative state. Fibroblast growth factor (FGF)–induced MAPK/ERK signaling pathway is an essential regulator of exit of embryonic stem cells from pluripotency^24,37–40^, with multiple lines of evidence suggesting that the pathway is under tight temporal control^25,41^. The regulation of endocytosis could provide a rapid and highly tunable means of regulating FGF signaling during these early stages. Going forward, it will be important to determine the link between miRNAs, endocytosis, cell signaling, and lineage diversification during this critical period of mammalian development.

## METHODS

### Cell culture and ESC monolayer differentiation

V6.5 ESCs were maintained on 0.2% gelatin-coated plates, cultured in Knockout DMEM (Invitrogen) supplemented with 15% FBS, LIF (1000U/mL), and when indicated, 2i (1uM MEK inhibitor PD0325901 and 3uM GSK3 inhibitor CHIR99021). Differential states of pluripotency from naïve to EpiLC transition were generated by removal of LIF and 2i as described in Krishnakumar et al., 2016. Briefly, ESCs were plated in ESC media. To initiate differentiation, LIF and 2i were removed approximately 24 hours after seeding. Cells were collected over indicated time points after removal of LIF and 2i.

### MiRNA mimic transfections

Mimic transfections were performed in indicated ESCs using miRIDIAN miRNA mimics (Dharmacon, ThermoFisher) using Dharmafect1 transfection reagent (Dharmacon, ThermoFisher) according to the manufacturers’ protocols. Control transfection to evaluate transfection efficiency were performed with every mimic transfection experiment using Dy547-labeled microRNA mimic (Dharmacon, ThermoFisher) and evaluated by flow cytometry.

### Generation of PFN2 ESC mutant lines by CRISPR-CAS9 gene editing

PFN2-deleted mutant Dgcr8-KO ESC lines were generated using clustered regularly interspaced short palindromic repeats (CRISPR) technology as described in Ran et al., 2013. Pairs of guide RNAs were designed against exon 2 of PFN2 using the CRISPR Design Tool (http://crispr.mit.edu/). Each guide RNA was cloned into a plasmid containing Cas9-GFP (OriGene plasmid GE100018, Rockville, MD). Paired guide-RNA plasmids were transfected into ESCs using FUGENE 6 (Roche) transfection reagent following the manufacturer’s protocol. The next day, GFP positive ESCs were sorted and plated at single-cell density onto gelatinized plates. Individual clones were genotyped for shifts in PCR product size in both alleles, and the resulting products were gel extracted and verified by Sanger sequencing. Positive clones were expanded and the absence of PFN2 protein was confirmed by western blot. PFN2 mutant Dgcr8-KO ESC lines with disruptions in the PFN2-Dynamin interaction site at PFN2 serine-138 (S138) were generated by single guide RNA designed against this site in exon 3 of PFN2, with processing as above. Guide sequences are listed in Supplementary Table 2.

### Endosomal uptake assays

pHrodo-conjugated transferrin and dextran are nonfluorescent at neutral pH and increasingly fluorescent with decreasing pH, with early endosomes labeled after 20 minutes and 5 minutes of uptake/internalization, respectively. Cells were serum-starved in HBSS + 1% BSA for 30 minutes at 37°C, then washed and resuspended with 150 μg/ml pHrodo Red conjugated transferrin or 80 μg/ml pHrodo Red conjugated 10000 MW dextran (Thermo Fisher Scientific) for 20 min at 4°C. Media was quickly added to the cells before incubation at 37°C, with one sample left at 4°C as an internal control. After indicated internalization times, cells were quickly moved back to 4°C, washed with cold PBS, and prepared for analysis by flow cytometry using the BD LSR II Flow Cytometer System (BD Biosciences).

### Immunoblot analysis

Cells were lysed with ice-cold RIPA lysis buffer containing EDTA-free protease inhibitor cocktail and PhosSTOP phosphatase inhibitor (Roche), with DNA-shearing by 21-gauge needle. Lysates were incubated at 4⍰°C for 15⍰min and centrifuged at 20,000g 4⍰°C on a table-top centrifuge. Protein was quantified using Bio-Rad protein assay (Bio-Rad). Thirty micrograms of protein was resolved on an SDS–PAGE gel. After electrophoresis, proteins were transferred onto Immobilon-FL (Millipore) PVDF membrane and processed for immunodetection. For immunodetection, immunoblots were incubated overnight at 4⍰°C with primary antibodies diluted in Intercept blocking solution (Licor) against alpha-tubulin at 1:3500 dilution (Sigma, T6074), profilin-2 at 1:700 dilution (BosterBio, PA2162), Phospho-p44/42 Erk1/2 Thr202/Tyr204 at 1:1000 dilution (Cell Signaling Technology, 4370), and p44/42 Erk1/2 at 1:1000 dilution (Cell Signaling Technology, 4696). Secondary infrared-dye antibodies were used at 1:10,000. Blots were scanned on a Licor Odyssey Scanner (Licor). Protein levels were quantified using ImageJ (http://rsb.info.nih.gov/ij/), and statistical calculations performed using at least 3 biological replicates.

### Quantitative real-time PCR

RNA for all qRT–PCR analyses was prepared using Trizol (Invitrogen) and quantified on a Nanodrop Spectrophotometer (ThermoFisher). RNA was DNase-treated using amplification grade DNaseI (Sigma). For qRT–PCR of mRNAs, DNase-treated samples were reverse-transcribed using the Maxima first-strand cDNA synthesis system (Thermo Scientific). Real-time quantitative PCR for mRNA was conducted with SYBR Green PCR master mix (Applied Biosystems) and performed on an ABI 7900HT (Applied Biosystems) according to the manufacturers’ protocols using primer sets listed in Supplementary Table 1.

### Growth rate assay

Counts were typically performed over three days after plating (on Day0) with Day1 counts and Day3 counts used to determine population growth rate. Growth rate was calculated using the equation [(log2(Yf) – log2(Yi)] / t where Yi is the initial count and Yf is the final count over a period of time t. Unless otherwise indicated, comparisons were performed relative to wild-type ESC cultured at the same time under the same culture conditions.

### Cell cycle analysis

Cell cycle phase distribution analysis was conducted using Click-iT EdU Alexa Fluor 647 Flow Cytometry Assay Kit (Life Technologies) and FxCycle Violet (Life Technologies) according to the manufacturer’s protocols. For EdU incorporation and DNA content analysis, cells were pulsed with 10⍰μM EdU (Life Technologies, Carlsbad, CA, USA) for 2 hours and subsequently stained according to manufacturer’s protocol. FxCycle Violet was added to click-treated cells before flow cytometry acquisition on an LSRII (BD Biosciences) with FACS Diva software (BD). A total of 10,000 events were collected for cell cycle analysis per sample. FlowJo software was used to process the data and determine the percentage of sample cells in each cell-cycle phase (FlowJo).

### FGF stimulation and ERK activation assay

Cells were pretreated with 3 μM BI-D1870 RSK inhibitor (Selleck Chemical) to potentiate FGF-stimulated ERK activation by blocking negative feedback regulation. For FGF-stimulation, cells were treated with 100ng/mL human recombinant FGF4 (R&D Systems, 235-F4) in the presence of 10 μg/mL heparin sulfate and 3 μM BI-D1870 RSK inhibitor at 37°C, then washed immediately with ice cold PBS and fixed with paraformaldehyde. Cells were permeabilized with 0.1% Triton-X and stained for intracellular phospho-p44/42 Erk1/2 Thr202/Tyr204 at 1:700 dilution (Cell Signaling Technology, 4370) and p44/42 Erk1/2 at 1:700 dilution (Cell Signaling Technology, 4696) for 30 minutes at 4°C followed by incubation with species-specific secondary antibodies Alexa Fluor 568 or 594 goat anti-mouse IgG at 1:200 dilution (Invitrogen, A11032) and Alexa Fluor 647 goat anti-rabbit IgG (Invitrogen, A21244) at 1:200 dilution for 30 minutes at 4°C. Cells were finally washed with cold PBS and prepared for analysis by flow cytometry using the BD LSR II Flow Cytometer System (BD Biosciences).

### Flow cytometry

Unless otherwise indicated, events were collected with FACS Diva software (BD) using a hierarchical gating strategy selecting for a homogeneous cell population (forward-scatter area vs. side-scatter area) of single cells (forward-scatter width vs. forward-scatter area). Dot plots and mean fluorescence intensities for gated events were generated using FlowJo analysis software (FlowJo).

### Statistical Analysis

Statistical details of experiments can be found in the figure legends, including the statistical test used, meaning of error bars, and p values. For all figures, the value of n indicates number of independent experiments and each dot indicates number of biological replicates (unless otherwise specified), defined as distinct dishes of cultured cells. Statistical analyses were performed using GraphPad Prism software.

## Data availability

The data that support the findings of this study are available from the corresponding author upon request.

## SUPPLEMENTARY FIGURES

**Supplementary Figure 1 (supplemental to Figure 1)**

(a) pH-labeled transferrin endosomal uptake assay in wild-type (WT), *Dgcr8*-KO, and *Dgcr8*-KO + miR-294 addback ESCs over indicated times, measured by change in mean fluorescence intensity (MFI) relative to control. *n* = 4 independent experiments. Error bars represent s.e.m. \**P* < 0.05, \*\*\*\**P* < 0.0001 (unpaired two-tailed t-test) (b-c) Sanger sequencing results showing mutations of the two biallelic mutant clones of (b) PFN2 exon 2 and (c) PFN2 exon 3 mutants used in experiments. (d) qRT-PCR analysis of PFN2 expression in wild-type (WT), *Dgcr8*-KO, PFN2 exon2 (PFN2-EX2KO) *Dgcr8*-KO mutants MT1 and MT2, and PFN2 exon3 (PFN2-S138) *Dgcr8*-KO mutants MT1 and MT2. *n* = at least 6 independent experiments. Error bars represent s.e.m. \*\*\*\**P* < 0.0001 (unpaired two-tailed t-test).

**Supplementary Figure 2 (supplemental to Figure 3)**

(a) Growth rates of indicated ESCs relative to wild-type (WT) ESCs under 1i (CHIR99021) vs. 2i (CHIR99021 + PD0325901 conditions. *n* = 3 independent experiments. Data shown as median ± quartiles. \*\*\**P* < 0.001, \*\**P* < 0.01 (unpaired two-tailed t-test). (b-d) Representative dot blots of source data for Fig 3e-g. (e) pH-labeled transferrin endosomal uptake assay in wild-type (WT) ESCs with and without added dynamin inhibition over indicated internalization times, measured by change in mean fluorescence intensity (MFI) relative to control. *n* = 4 independent experiments. Error bars represent s.e.m. \*\*\*\**P* < 0.0001 (unpaired two-tailed t-test)

**Supplementary Figure 3 (supplemental to Figure 4)**

(a) qRT-PCR analysis of Lats2 and Cdkn1a expression in wild-type (WT), *Dgcr8*-KO, *Dgcr8*-KO with addback of miR-294 mimic (+294) or PFN2-depletion (PFN2-EX2KO) ESCs under naïve conditions n=3 independent experiments. Error bars represent s.e.m. \*\*\**P* < 0.001, \*\**P* < 0.01, \**P* < 0.05 (unpaired two-tailed t-test). (b) qRT-PCR analysis of EpiLC markers *Otx2* and *Dnmt3b* expression at indicated time points over ESC to EpiLC transition in wild-type (WT), *Dgcr8*-KO, *Dgcr8*-KO with addback of miR-294 mimic (+294) or *Dgcr8*-KO; PFN2-EX2 DKO ESCs n=3 independent experiments. Error bars represent s.e.m. \*\*\*\**P* < 0.0001 (unpaired two-tailed t-test).

**Supplementary Figure 4 (supplemental to Figure 5)**

(a) Figure exemplifying the gating strategies used. Briefly, viable cells were selected and cell debris excluded using the FSC vs SSC gate. We selected for singlets using FSC-width vs FSC-area gates. (b) Unstained cells of the same cell line or parental cell line were used as general negative controls for fluorescence. In the case of dual-color analysis, single-color controls were used to determine the boundaries between negative and single-color positive and dual-color positive cell populations. (c) Representative dot blots of source data for one replicate of ERK activation time course experiments summarized in Fig 5i.

**Supplementary Figure 5**

Unprocessed western blots from indicated figures.

