## Supplementary figures and images for "MicroRNA-dependent inhibition of PFN2 orchestrates ERK activation and pluripotent state transitions by regulating endocytosis"

### Supplemental Figures

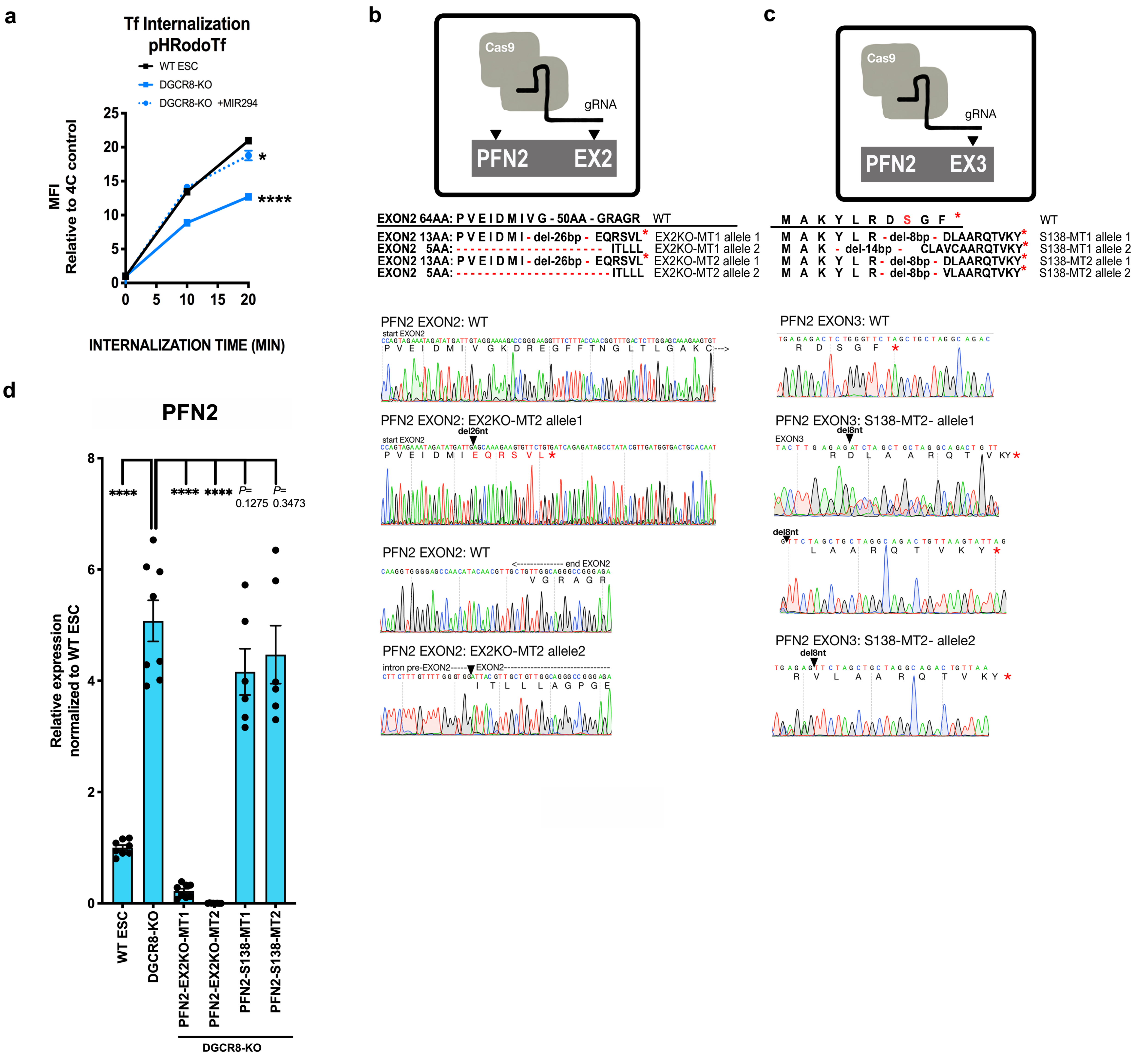

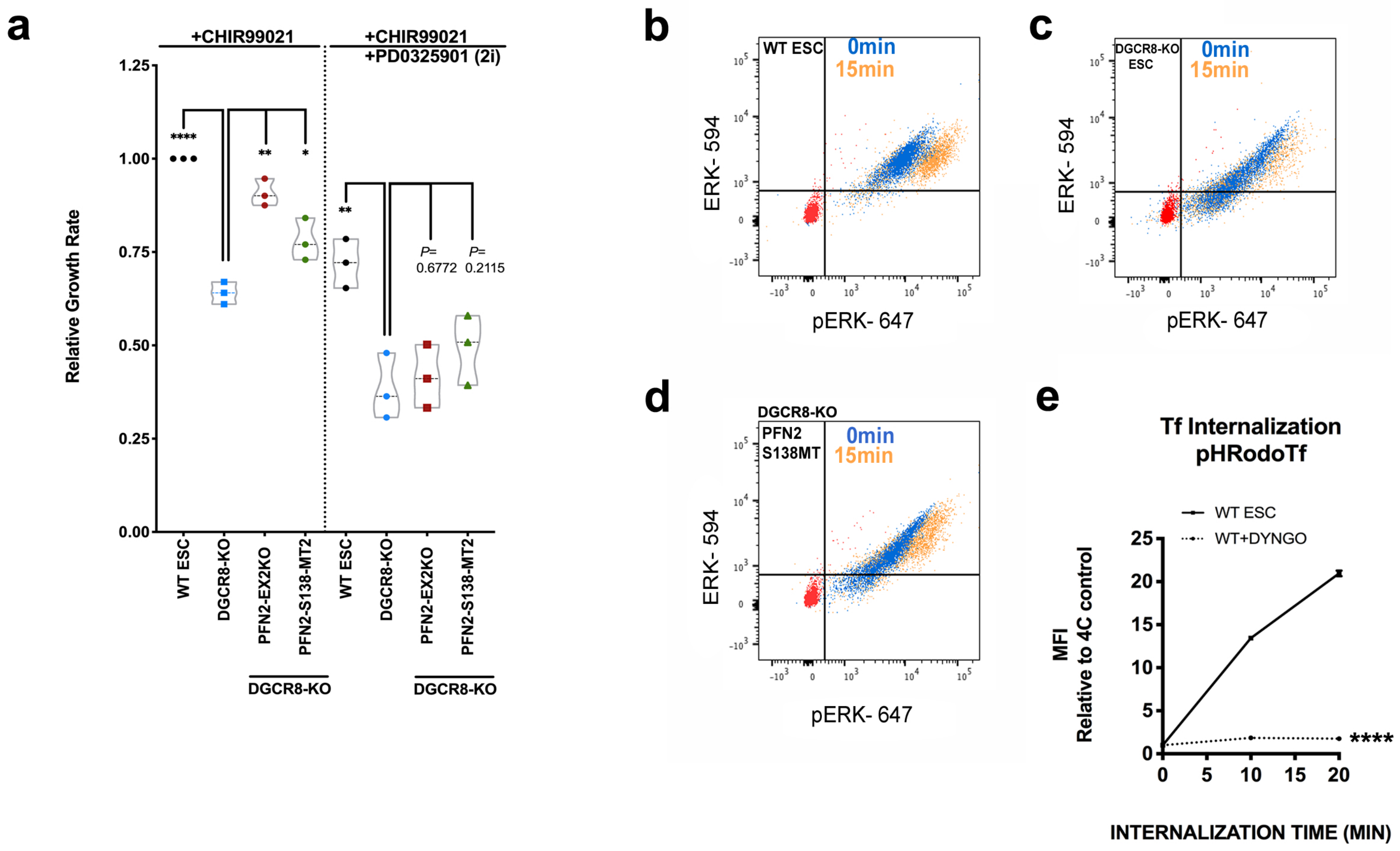

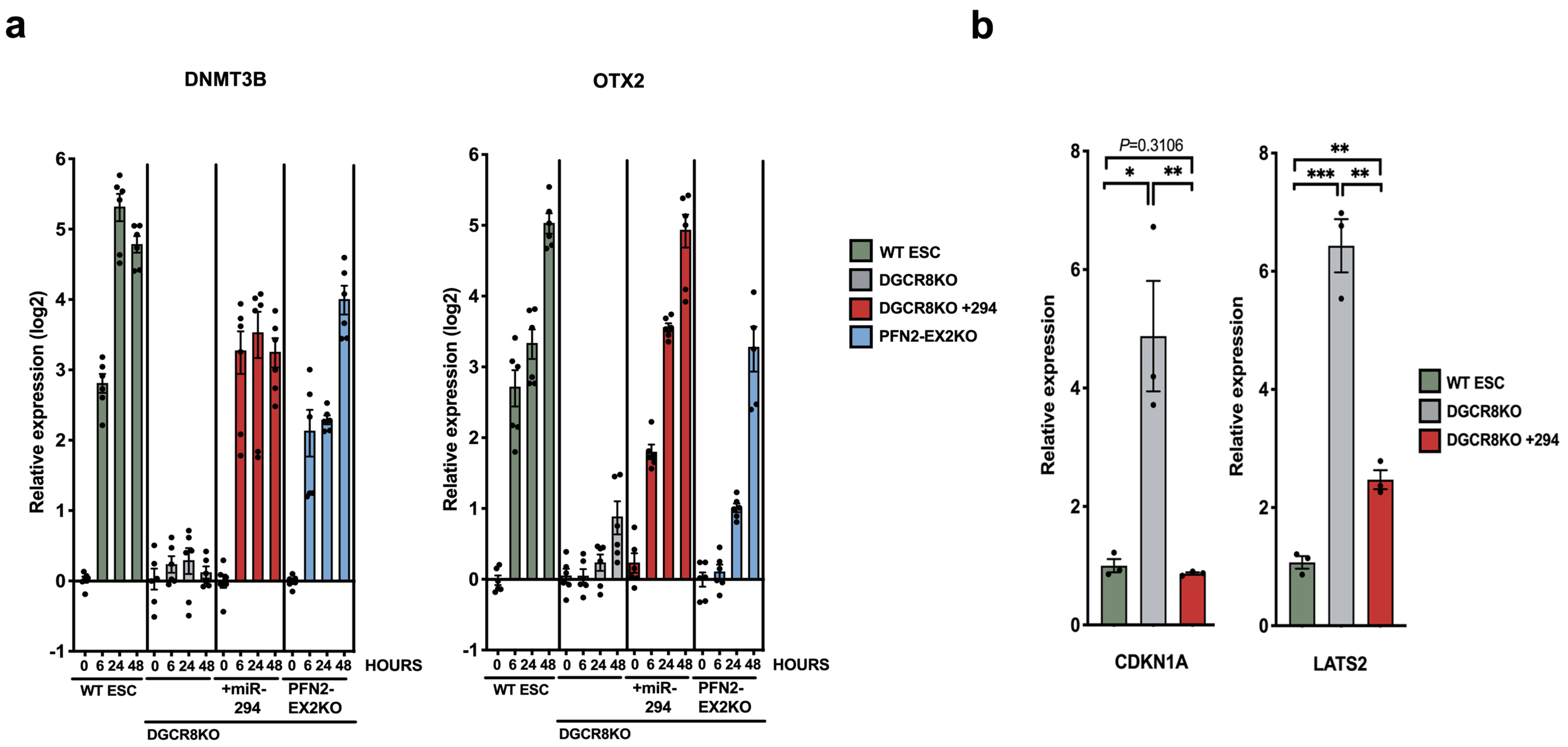

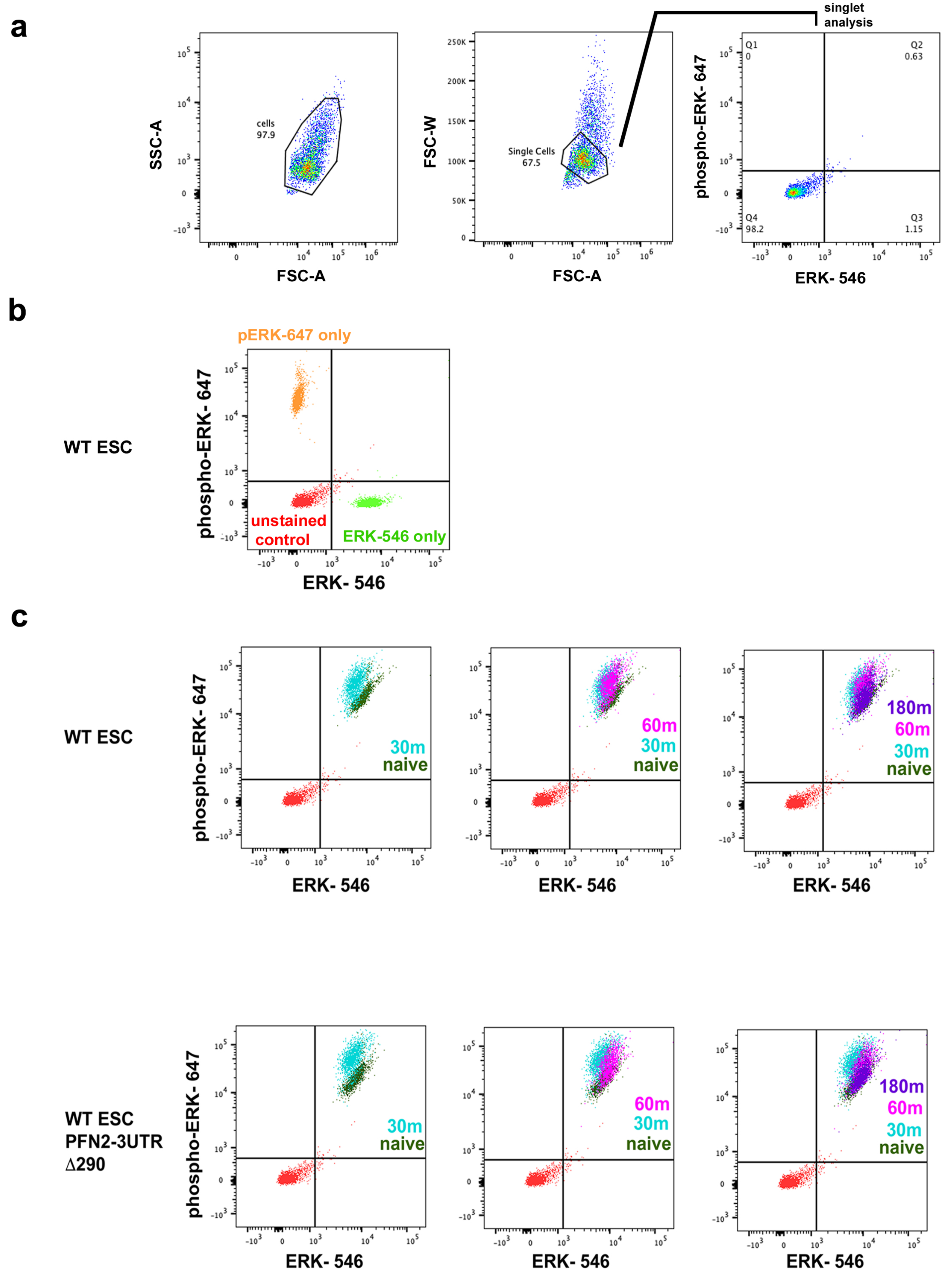

#### Figure S5

Fig. 1a

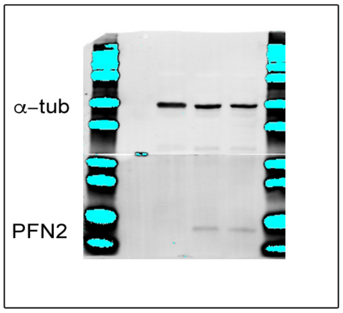


Fig. 1f

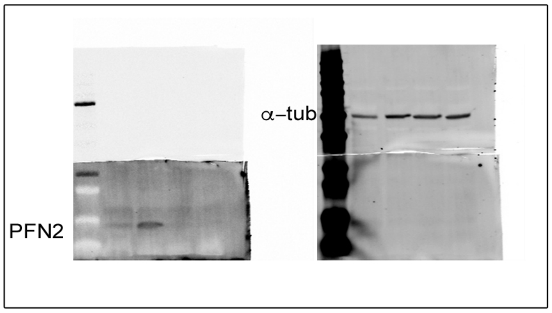


Fig. 1g

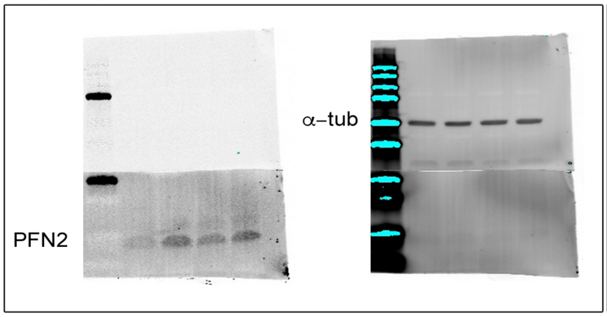


Fig. 3a

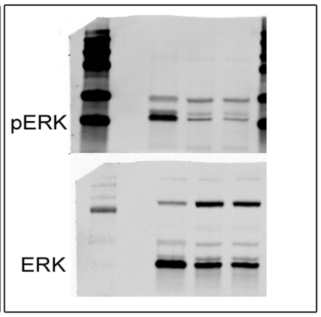


Fig. 3b

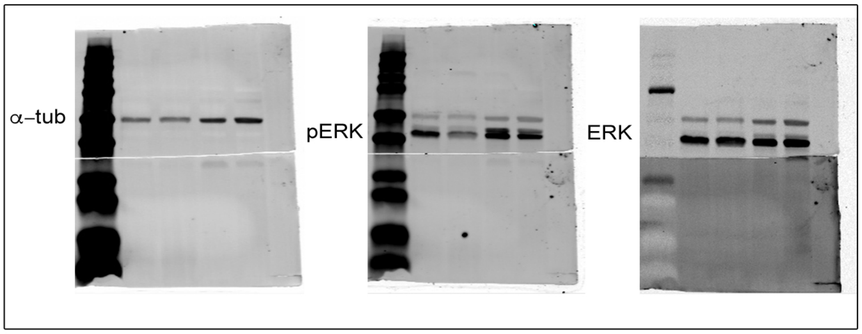


Fig. 3c

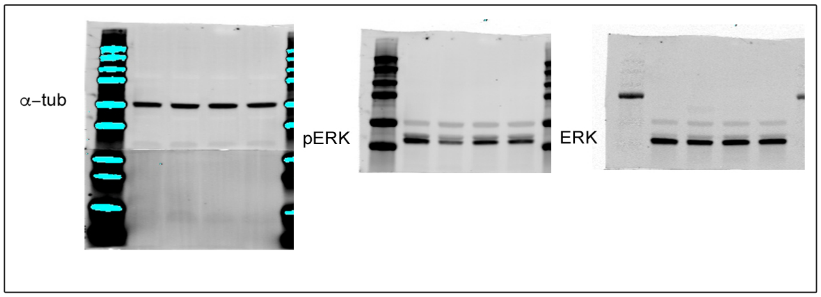


Fig. 5h

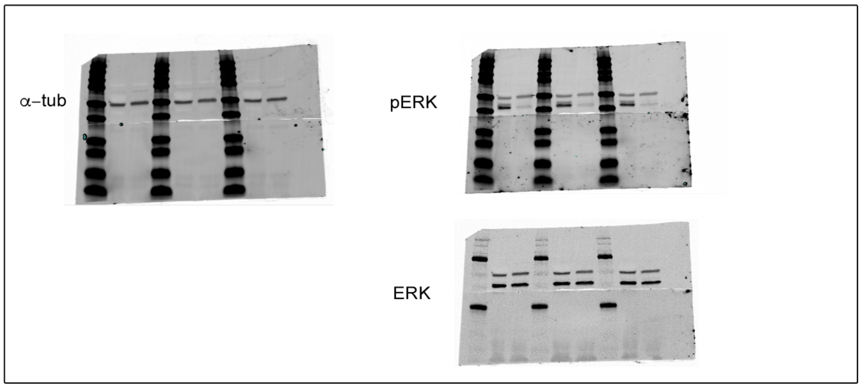
